# Calcyphosine is a microtubule-associated protein required for spindle formation and function

**DOI:** 10.1101/2023.12.29.573632

**Authors:** Bipul Setu, Qian Nie, Grace Echele, Susan A. Spencer

**Affiliations:** Department of Biology, Saint Louis University, 3507 Laclede Ave., St. Louis, MO 63103; Division of Infectious Diseases, Department of Internal Medicine, Washington University School of Medicine, 660 Euclid Ave., St. Louis, MO 63110; Belay Diagnostics, 1375 W Fulton St, Chicago, IL 60607

## Abstract

Calcyphosine (CAPS) is a highly conserved but little explored calcium-binding protein that shows elevated expression in many forms of human cancer. Here we uncover a role for CAPS in spindle formation during mitosis. Our experiments suggest that CAPS is a microtubule-binding, spindle-associated protein that helps create the kinetochore fibers that bind and segregate chromosomes. Knockdown of CAPS causes a variety of defects during mitosis, including uncongressed chromosomes and multi-polar spindles, as well as high levels of apoptosis and a reduced mitotic index. We find that CAPS promotes microtubule bundling, both in vitro and in cells, and knockdown of CAPS leads to reduction of thick k-fibers in the mitotic spindle. The high level of CAPS observed in many forms of cancer suggests that CAPS may promote cell proliferation, but our results indicate that CAPS overexpression has little effect on the cell cycle. This suggests that the high level of CAPS expression may be a consequence of cancer, rather than a driving force for cell proliferation.

## Introduction

Calcyphosine (CAPS) is a small, highly-conserved calcium-binding protein of unknown function (Yuasa et al., 2002). It was originally identified as a target for phosphorylation by protein kinase A downstream of TSH signaling (Lecocq et al., 1979; Lecocq et al., 1990) though that characterization was later qualified (El Housni et al., 1997). In recent years, proteomic screens have implicated high levels of CAPS in a variety of human cancers, including endometrial cancer, colorectal cancer, and esophageal squamous cell carcinoma, suggesting a possible role for CAPS in cell cycle regulation (de Bont et al., 2007; Li et al., 2010; Shao et al., 2016; Li et al., 2017). Yet despite these studies, little is known about the function of CAPS or whether it plays a role in cancer progression.

The crystal structure of CAPS has been solved and suggests that when bound to calcium, CAPS undergoes a structural shift that regulates its binding to hydrophobic phenyl-Sepharose (Oyama et al., 1997; Dong et al., 2008). This indicates that calcium may act as a switch that regulates the binding of CAPS to other proteins. From both its amino acid sequence and its crystal structure, CAPS bears similarities to calmodulin, a calcium-binding protein whose interactions with enzymes such as myosin light-chain kinase, calcineurin, and CaM-kinase II are modulated by intracellular calcium concentrations (Andrews et al., 2020). While binding partners for CAPS have not yet been verified, high throughput proteomic screens suggest that CAPS coprecipitates tubulin, the zinc-finger protein ZC2HC1A, and the microtubule spindle-associated protein Hepatoma Upregulated Protein (HURP) (Huttlin et al., 2021). The inclusion of both tubulin and HURP on this short list suggests that CAPS may associate with the microtubule mitotic spindle.

Many microtubule-associated proteins influence microtubule polymerization and stability throughout the cell (Brouhard and Rice, 2018; Lawrence et al., 2023). Formation of the mitotic spindle additionally requires clustering of some microtubules into bundles (reviewed in Valdez et al., 2023). At least two classes of microtubules undergo bundling: kinetochore fibers (k-fibers), which form parallel bundles that attach to the kinetochores of chromosomes, and bridging fibers, which form anti-parallel bundles that cross-link other k-fibers near the metaphase plate and are important for spindle architecture and mechanics. K-fibers are groups of 9-20 individual microtubule strands that are linked to the centrosome and associate with one another through bridges composed of proteins such as HURP, ch-TOG, TACC, and clathrin (Sillje et al., 2006; Koffa et al., 2006; Wong and Fang, 2006; Booth et al., 2011). Bridging fibers, on the other hand, require the protein PRC1 for their association, and are found near the metaphase plate, where they link and organize k-fibers attached to sister chromatids (Subramanian et al., 2010; Kajtez et al., 2016; Simunic and Tolic, 2016; Polak et al., 2017; Matkovic et al., 2022). The distinct protein components and localization of these microtubules bundles distinguish them from one another in the mitotic spindle.

CAPS conformation appears to be regulated by calcium-binding. Calcium is required at several stages for cells to progress through mitosis (Tombes and Borisy, 1989). Increased intracellular calcium appears to speed the onset of anaphase from metaphase (Izant, 1983), while chelation of calcium inhibits cell cycle progression. BAPTA-AM, a cell-permeable calcium chelator, produces defects in mitotic spindle organization and CENPF localization, though some of BAPTA-AM’s effects on microtubules and other cytoskeletal elements may be unrelated to calcium chelation (Xu et al., 2003; Phengchat et al., 2017; Saoudi et al., 2004; Furuta et al., 2009). Other methods of reducing calcium levels, however, produce similar conclusions: chelation of nuclear calcium, for instance, by specific targeting of calcium-binding proteins, also blocks cell cycle progression (Rodrigues et al., 2007). Calcium levels are locally high near spindle poles; this is thought to derive from IP3-mediated release of calcium from nearby ER and correlates with the centrosomal localization of calmodulin (Erent et al., 1999; Moisoi et al., 2002). When calcium is reduced specifically at spindle poles, progression through cell division is inhibited (Helassa et al., 2019). Taken together, these results suggest that calcium is locally required in the nucleus and at spindle poles for cells to progress through mitosis.

Here we present evidence that CAPS binds tubulin in the presence of calcium and associates with the mitotic spindle in dividing cells. We find that reduction of CAPS produces spindle defects, uncongressed chromosomes, and increased cell death. Consistent with this, CAPS appears to promote microtubule bundling into kinetochore fibers (k-fibers) and satisfaction of the spindle assembly checkpoint (SAC). Finally, given the high level of CAPS present in many cancers, we examine the effects of CAPS overexpression. Our results suggest that CAPS promotes formation of k-fibers and that its absence leads to defects in spindle structure and chromosome congression.

## Results

### CAPS is a microtubule-binding protein

Based on its amino acid sequence, CAPS appears to be a human ortholog of the *Drosophila* protein CG10126. Previous work in our lab has indicated that CG10126 binds robustly to tubulin in a calcium-dependent manner (*in preparation*); consistent with this, recent high throughput screens also suggest that CAPS is a tubulin binding protein (Huttlin et al., 2021). To confirm this interaction, we created a FLAG-epitope-tagged form of CAPS and transfected it into cultured HEK293 cells. Lysates from these cells were used for immuneprecipitations with anti-FLAG Sepharose in the presence of calcium or the calcium-chelator EGTA. Our results indicate that CAPS co-precipitates tubulin robustly in the presence, but not the absence, of calcium (Fig. 1A).

**Figure 1:**
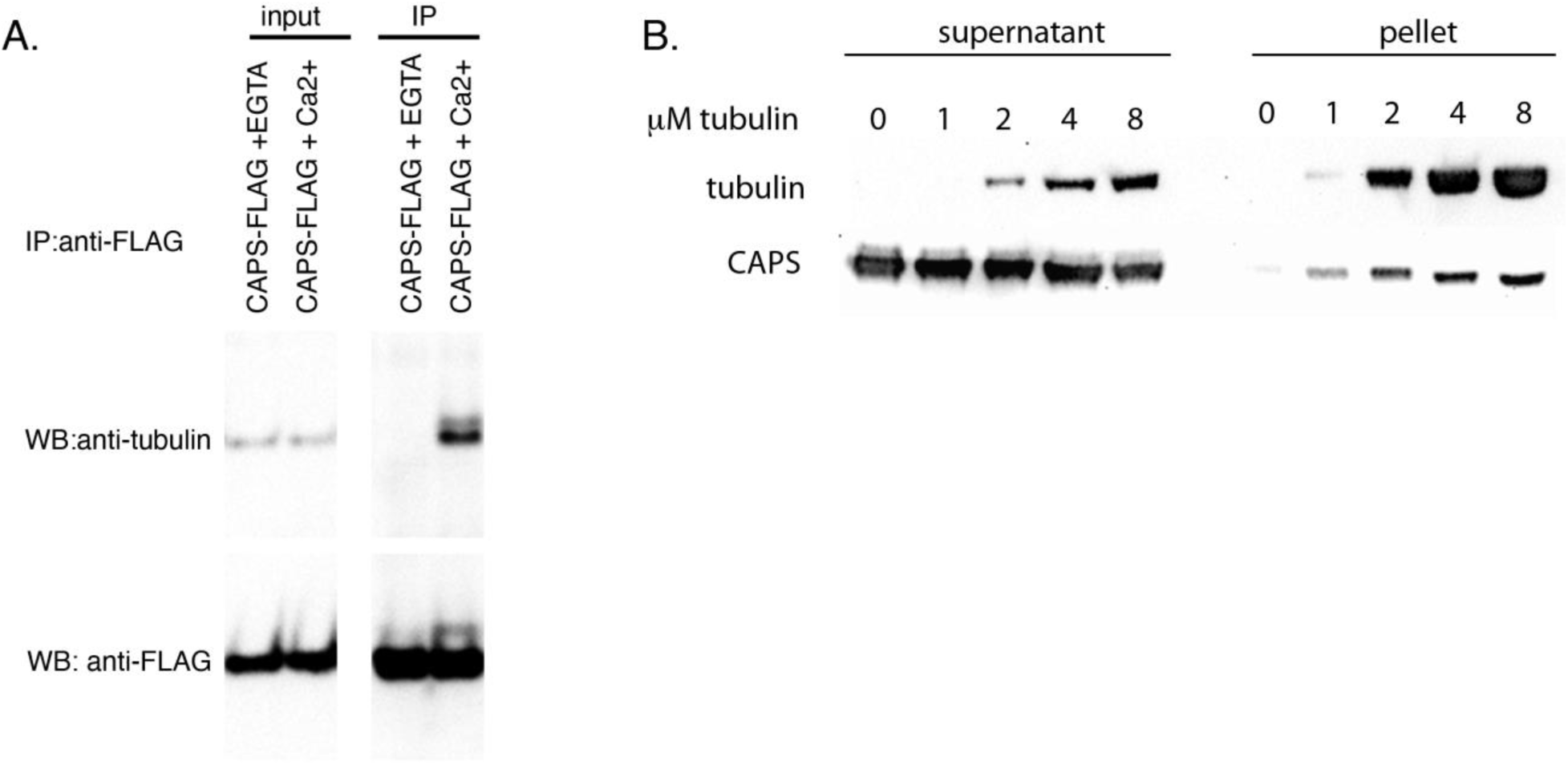
CAPS co-precipitates with tubulin and microtubules in the presence of calcium. A) Western blot shows immuneprecipitations of CAPS-FLAG from transfected HEK293 in the presence of EGTA or calcium. CAPS only coprecipitates tubulin in the presence of calcium. B) Western blot of spin down assay. Purified and polymerized microtubules were centrifuged through a glycerol cushion in the presence of 50uM calcium; polymerized microtubules and bound protein are in the pellet. CAPS in the pellet increases with greater microtubule concentrations.

Since cold depolymerizes most intact microtubules (Li and Moore, 2020) and our immuneprecipitations were performed on ice, our results suggest that CAPS can bind tubulin dimers but do not indicate whether CAPS can also bind to polymerized microtubules. To test this, we performed a microtubule spin-down assay: polymerized microtubules (and any associated proteins) were spun at high speed in the presence of calcium through a glycerol cushion to form a pellet, while the supernatant retained tubulin dimers and unbound proteins. CAPS coprecipitated with microtubules, and the amount of CAPS in the pellet rose with increasing microtubule concentrations, suggesting that CAPS is capable of binding to polymerized microtubules as well as individual dimer subunits (Fig. 1B).

### CAPS is associated with central spindle microtubules

These findings suggested that CAPS might localize to the microtubule cytoskeleton in cells. Further, Huttlin et al. 2021 provides evidence that CAPS binds DLGAP5/HURP, a known component of the mitotic spindle (Tsou et al., 2003; Sillje et al., 2006; Wong and Fang, 2006; Koffa et al., 2006; Zhang et al., 2018), raising the possibility that CAPS might be spindle-associated. To observe CAPS localization, we used a commercially available antibody against CAPS for immunocytochemistry in human cell lines. SK-BR-3, a human breast cancer cell line that has a high level of CAPS expression and U251, a glioblastoma cell line that has moderate expression of CAPS, were chosen for analysis (Karlsson et al., 2021; Human Protein Atlas, proteinatlas.org). In mitotic cells of both lines, CAPS appears concentrated along the spindle at all stages (Fig. 2A, Supplemental Fig. 1). Similar staining patterns were observed with a CAPS construct containing an N-terminal gfp tag (Fig. 2B). Interestingly, CAPS appears to localize only to central microtubules of the spindle, which include bipolar microtubules and kinetochore fibers, but not to astral microtubules (Fig. 2C, C’). Consistent with this, in interphase cells, CAPS has a punctate appearance and is distributed between the cytosol and nucleus, but does not appear to be strongly associated with cytosolic microtubules (Fig. 2D, D’). These results suggest that while CAPS can bind polymerized microtubules, it is not associated with all microtubules in the cell, but instead specifically binds to the central microtubules of the mitotic spindle.

**Figure 2:**
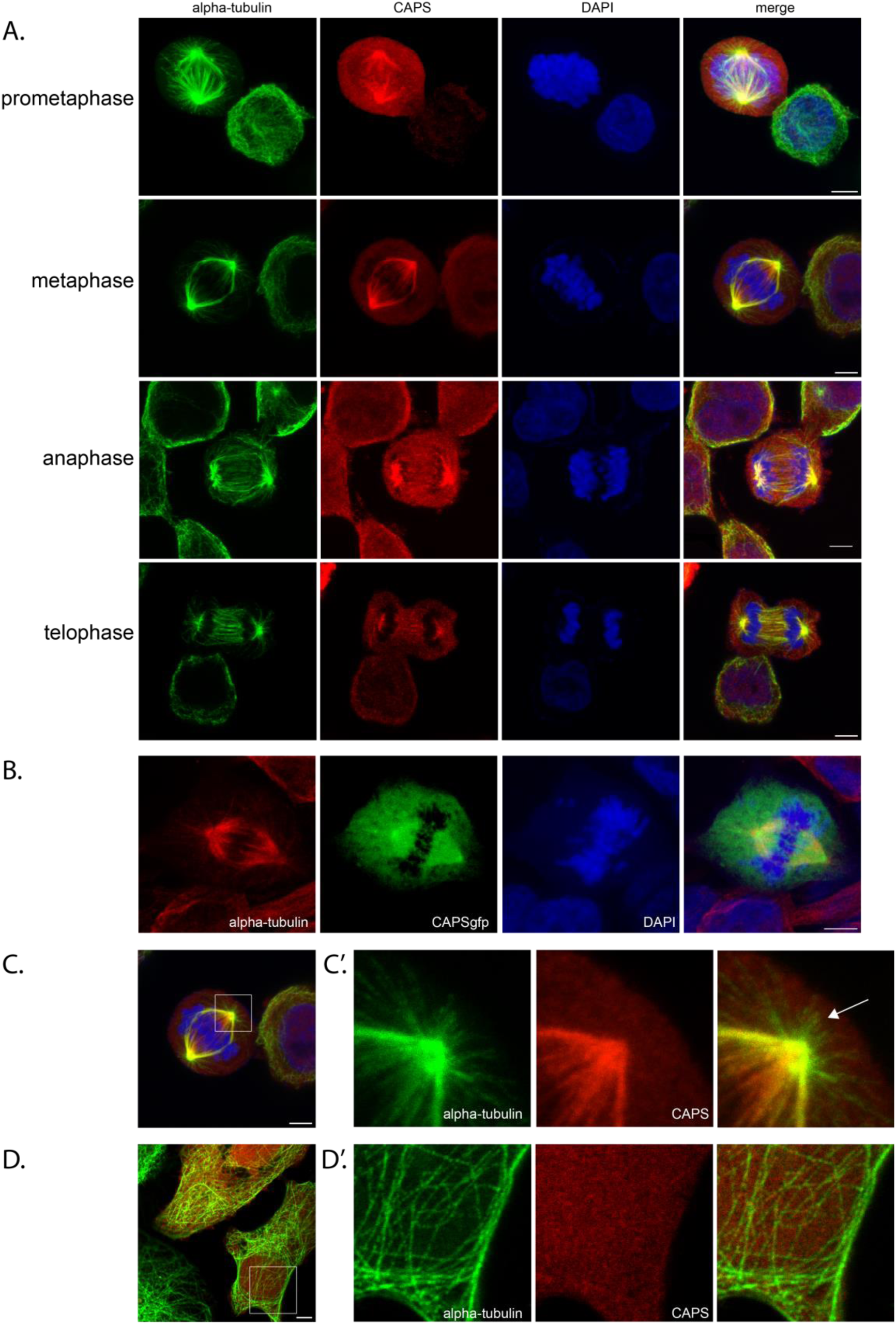
CAPS is associated with the mitotic spindle A. Cultured human SKBR3 cells were fixed and stained with antibodies against CAPS (red) and alpha-tubulin (green); DAPI is shown in blue. Fluorescent immunostaining shows CAPS along microtubules between the spindle poles at all stages of mitosis. B) U251-MG cells transfected with CAPS-gfp show a pattern of CAPS localization similar to that seen with CAPS antibody. C) Metaphase cell from (A) showing blow up of area in square (C’). The localization of CAPS to the spindle is limited to central microtubules; CAPS is not found along astral microtubules (arrow). D. In interphase cells, CAPS does not appear to colocalize with cytoplasmic microtubules; D’ blow up of square in D. Staining as in (A). Images are max Z-projections of confocal stacks. Scale bars are 5um in all pictures.

### Loss of CAPS leads to spindle defects, reduced mitotic index, and increased cell death

The spindle-specific localization of CAPS raised the possibility that it might play a role in spindle assembly or stability. To test this, we reduced the level of CAPS by transfecting U251 cells with a CAPS-specific siRNA. As seen in Figure 3A, introduction of siRNA reduced the level of CAPS in U251 cells by ~77%; this siRNA was then used to assess phenotypes associated with CAPS knockdown. Because the sequence of this siRNA is complementary to sequences in the 3’ UTR of CAPS mRNA, it was also possible in many experiments to transfect cells with a cmv-CAPS-FLAG overexpression construct to ensure that phenotypes observed were specifically due to CAPS knockdown.

**Figure 3:**
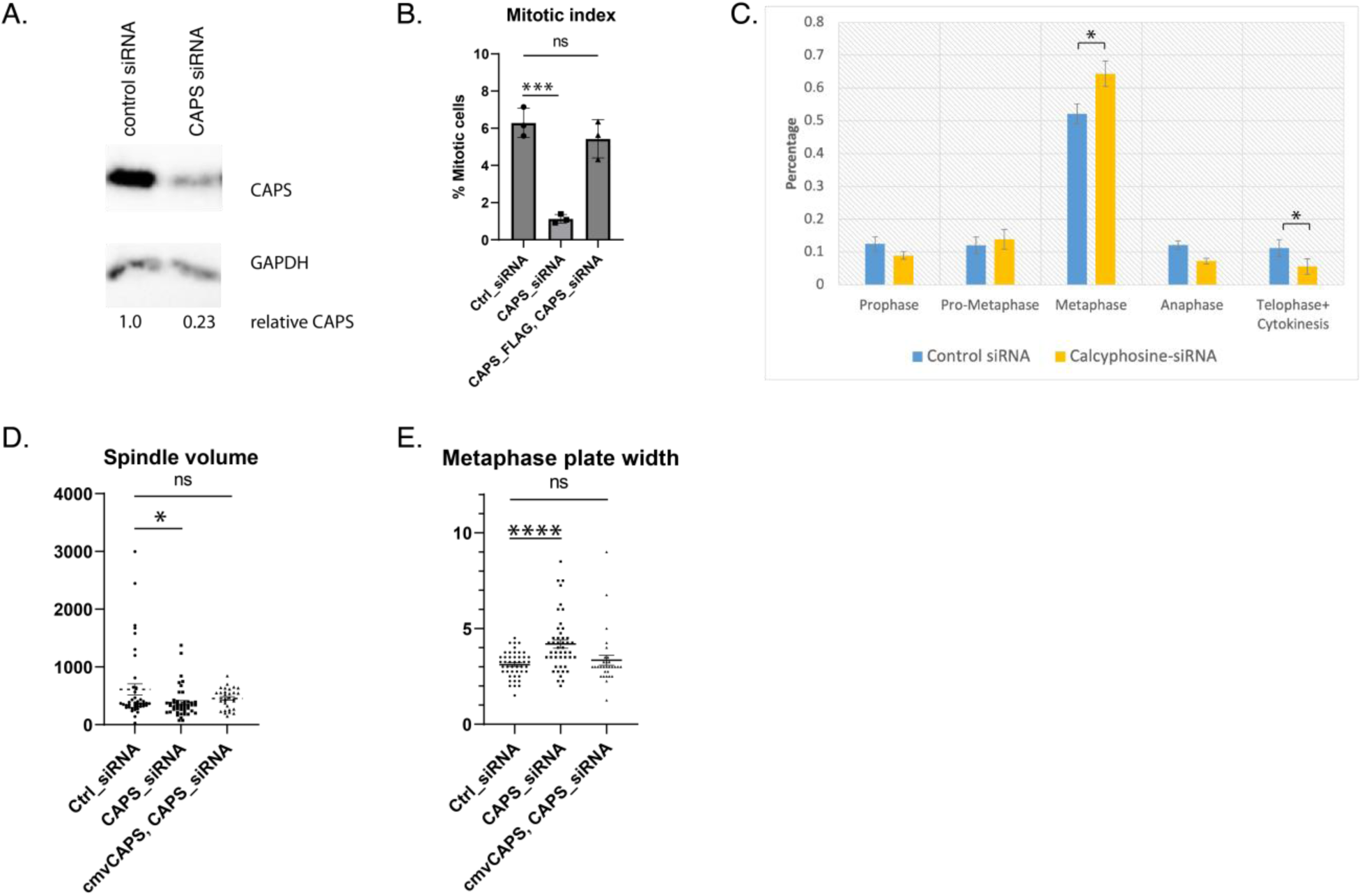
CAPS knockdown in U251MG cells leads to changes in mitosis. A. Western blot with anti-CAPS antibody showing that transfection of CAPS siRNA reduces CAPS protein level by approximately 77%. B. Knockdown of CAPS reduces the mitotic index of U251MG cells from 6.3% to 1.1% (p=.004); simultaneous expression of a CAPS rescue construct restores the mitotic index to 5.4% (20 fields of cells counted for each condition; three independent experiments). C. Knockdown of CAPS increases the percentage of cells in metaphase (from 54% to 64%; p=.01), while slightly decreasing the percentage of cells in telophase (from 11% to 5.6%; p=.01). (50 mitotic cells evaluated/trial; N of 3). D) and E) The Spindle 3-D Image J plugin (Kletter et al., 2022) was used to evaluate characteristics of the mitotic spindle in cells transfected with control siRNA, CAPS siRNA, or CAPS siRNA plus CAPS-FLAG (rescue). D. CAPS knockdown leads to a decrease in spindle volume (610um^3^ for control siRNA, 380 um^3^ for CAPS siRNA; p=.02). E. CAPS knockdown also leads to an increase in the metaphase plate width (width of chromosomes on the equator; 3.1 um for control siRNA vs. 4.2 for CAPS siRNA; p<0.01. For D and E, 35-50 cells were evaluated per condition with pooled data from three trials.

To test the role of CAPS in spindle assembly, control and CAPS-depleted cells were grown on coverslips and the mitotic spindle was visualized via immunofluorescence with antibodies to alpha-tubulin and gamma-tubulin; chromosomes were stained with DAPI. Cells with reduced CAPS had a lower mitotic index than controls (3B, 1.1% vs. 6.3%, p-value = <.01), suggesting that loss of CAPS leads to defects that limit entry into mitosis. Among mitotic cells, knockdown of CAPS slightly increased the proportion of cells in metaphase, indicating that CAPS loss might block entry to anaphase (Fig. 3C). We further analyzed cells with bipolar metaphase spindles using the Spindle 3D plugin in Fiji (Fig. 3D, E); (Kletter et al., 2022); 40-50 cells from 3 independent experiments were included in the calculations. This analysis showed no significant difference between control siRNA and CAPS siRNA treatment in spindle length (10.07 vs. 9.65um, p>.05; not shown). However, spindle volume was reduced with CAPS knockdown (610 um3 vs. 380 um3; p=.02; Fig. 3D, and there was a significant difference in the width of chromosomes along the metaphase plate between control cells and those with reduced CAPS (3.12 um vs. 4.19um; p=.0001; Fig. 3E). Overexpression of CAPS reverted the spindle volume and chromosome width along the metaphase plate to control levels, indicating that these alterations were due to loss of CAPS. Together, these results suggest that while CAPS is not essential for the formation of the mitotic spindle, its loss leads to a decrease in spindle volume, disorganization in the alignment of chromosomes along the metaphase plate, and ultimately, to a prolonged metaphase period and decrease in the number of mitotic cells.

The usual narrow alignment of chromosomes along the metaphase plate is induced by attachment of chromosome kinetochores to kinetochore fibers (k-fibers) on either side of the mitotic spindle (Tanaka and Desai, 2008). The increased width of chromosomes along the metaphase plate observed with CAPS knockdown suggested that there might be defects in this attachment. Indeed, we found that loss of CAPS led to a dramatic increase in the incidence of metaphase cells with uncongressed chromosomes compared to controls (4A, B; 46% vs. 9%, p-value<.01). Multipolar spindles sometimes arise when anaphase is delayed due to activation of the spindle assembly checkpoint (SAC) (Maiato and Logarinho, 2014). Consistent with this, there was also a significant increase in the incidence of multipolar spindles with CAPS knockdown (Fig. 4C, D; 16% vs. 5%, p-value=.03). Overexpression of CAPS in siRNA-treated cells rescued these defects. To test if high levels of SAC proteins were associated with the uncongressed chromosomes in our siRNA knockdowns, we stained cells with antibodies against HEC1, which marks the kinetochore, and BubR1, a protein retained at centromeres of unattached chromosomes. We found that the BubR1:HEC1 ratio is substantially higher on these chromosomes, consistent with a loss of attachment between these chromosomes and kinetochore microtubules (Fig. 4E).

**Figure 4:**
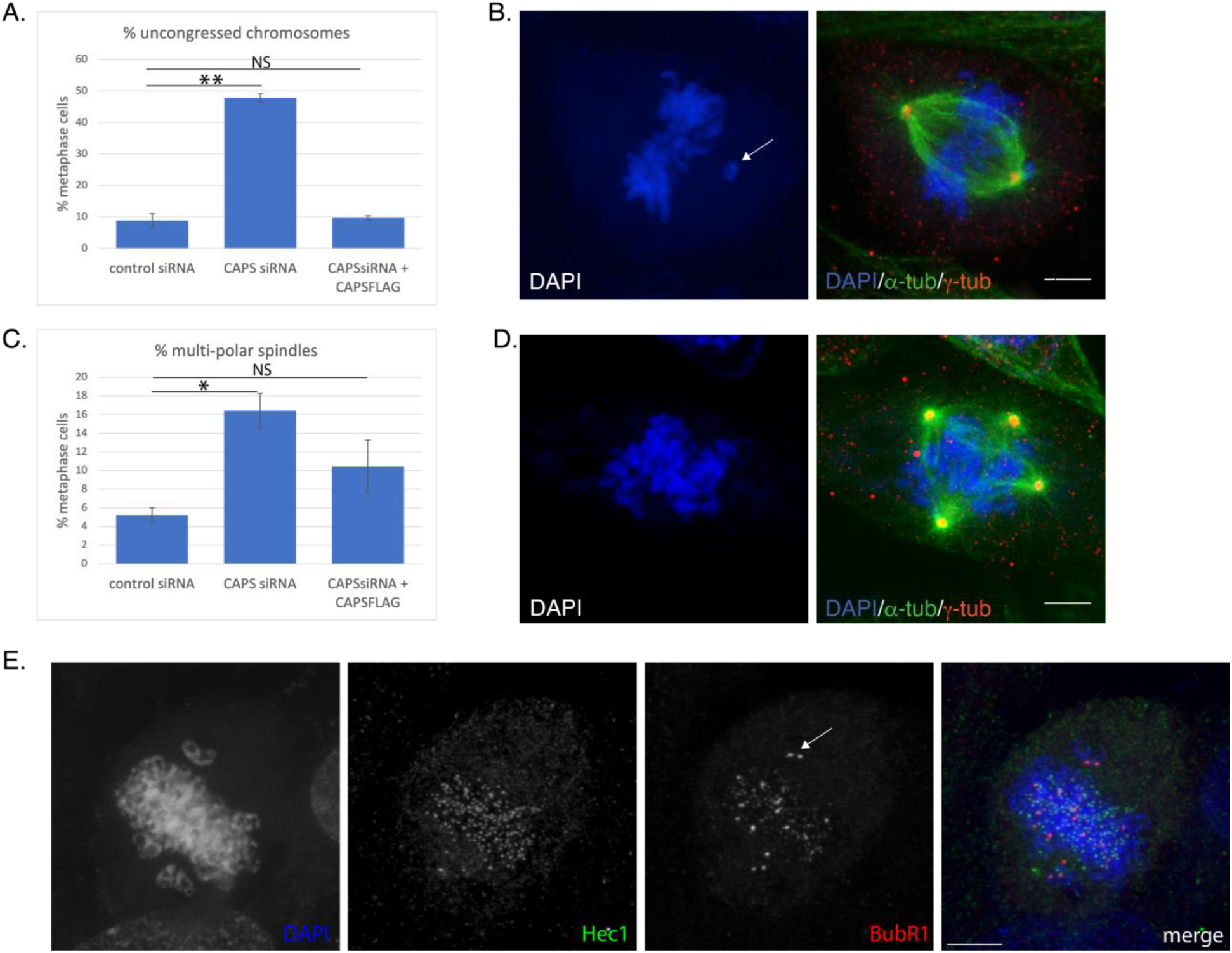
Knockdown of CAPS alters spindle morphology and chromosome congression. A) and B) Fluorescent immunohistochemistry and confocal microscopy of cells transfected with control siRNA, CAPS siRNA, or CAPS siRNA plus CAPS-FLAG (rescue) was used to evaluate the percentage of mitotic cells containing uncongressed chromosomes (arrow in B) (8.8% for control siRNA, 47.8% for CAPS siRNA; 9.7% for CAPS siRNA + CAPS-FLAG; Fisher’s exact test p<.01; 44 cells per condition in each of three independent experiments. C) and D) Mitotic cells were evaluated for the presence of multipolar spindles as in A), using gamma-tubulin as a marker for spindle poles. CAPS kd led to increased presence of multipolar spindles; Fisher’s exact test, p=.03; 44 cells per condition in each of three independent experiments. E) Cells transfected with CAPS siRNA were also evaluated for the presence of the spindle assembly checkpoint protein BubR1; Hec1 is present on all kinetochores, while BubR1 is elevated on uncongressed chromosomes (arrow) and some chromosomes along the metaphase plate.

Despite the maintenance of the SAC in some CAPS-knockdown cells, we observed that many cells would proceed through mitosis, often creating multipolar spindles before anaphase occurred. This resulted in production of 3-4 daughter cells with abnormal numbers of chromosomes. (Movies S2a and S2b). This surprising procession through anaphase in the presence of an activated SAC was also observed by Wong and Fang, 2006 with HURP knockdown. Consistent with this phenomenon, cells treated with CAPS siRNAs showed increased levels of apoptosis, as evidenced by staining cells with antibodies specific for cleaved Caspase 3: 7.1% of CAPS siRNA-treated cells contained activated Caspase-3, compared to 0.2% of control cells (Fig. 5A,B, p=.0001). Similarly, incorporation of Bromodeoxy-uridine (BrdU) at exposed DNA 3’ ends was much higher with CAPS knockdown than in controls (10.1% in CAPS knockdown vs. 3.0% in controls, p=.0001; Fig. 5C,D). Together, these results suggest that CAPS knockdown produces a loss of chromosome congression during metaphase that leads to uneven distribution of chromosomes and eventual cell death.

**Figure 5.**
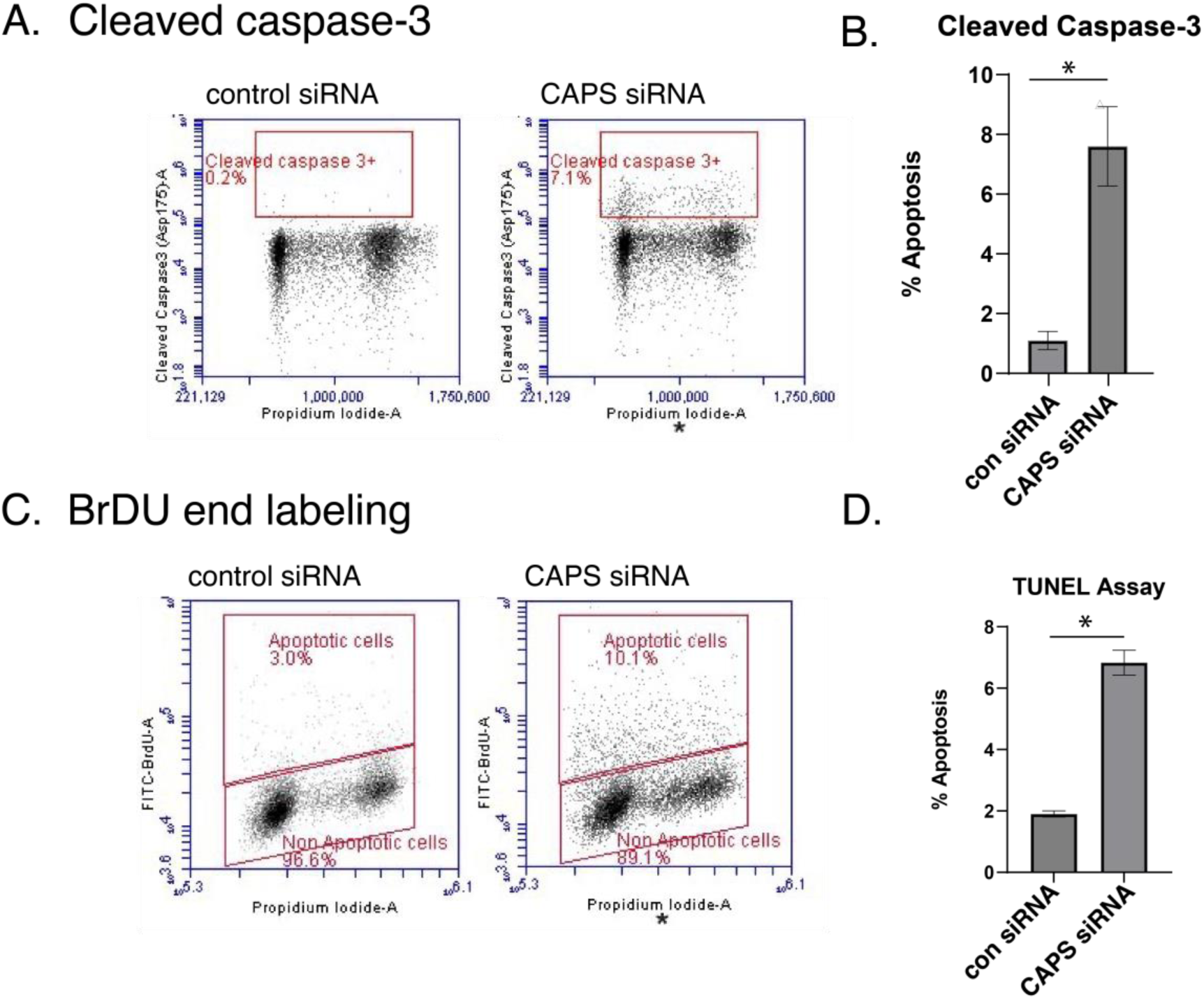
Knock down of CAPS induces apoptosis. A, B. Apoptosis in CAPS or control siRNA-treated cells was analyzed by staining with antibodies against cleaved Caspase 3 and flow cytometry. CAPS knockdown increased the percentage of cells containing cleaved caspase 3 from 0.62% to 7.9% (p value=.0001) C.D. CAPS or control siRNA-treated cells were analyzed using TUNEL (BrDU-labeling of exposed 3’ DNA ends) to identify apoptotic cells. CAPS knockdown increased the apoptotic cell population from 3.0% to 10.1% (p value=.0001)

### Loss of CAPS is associated with reduced k-fiber formation

K-fibers are bundles of ~9-20 microtubules linked by an intermicrotubule mesh (McEwen et al., 1997; O’Toole et al., 2020; Kiewisz et al., 2022); this bundling is necessary for attachment of k-fibers to chromosome kinetochores. The wider metaphase plate and loss of chromosome congression that we observed with CAPS knockdown, together with the localization of CAPS along central spindle microtubules, suggest that CAPS may contribute to bundling of microtubules into k-fibers. In confocal micrographs of CAPS-depleted cells, we observed spindle microtubules often appeared to be both less robust and less grouped than those of controls (6A, A’). To observe this more closely, we used expansion microscopy to gain a higher resolution view of the spindle in control and CAPS knockdown cells. In controls, thick k-fibers are readily apparent, while in many CAPS knockdown cells, central spindle microtubules appear to be primarily narrow strands, suggesting a loss of microtubule bundling (Fig. 6B, B’).

**Figure 6.**
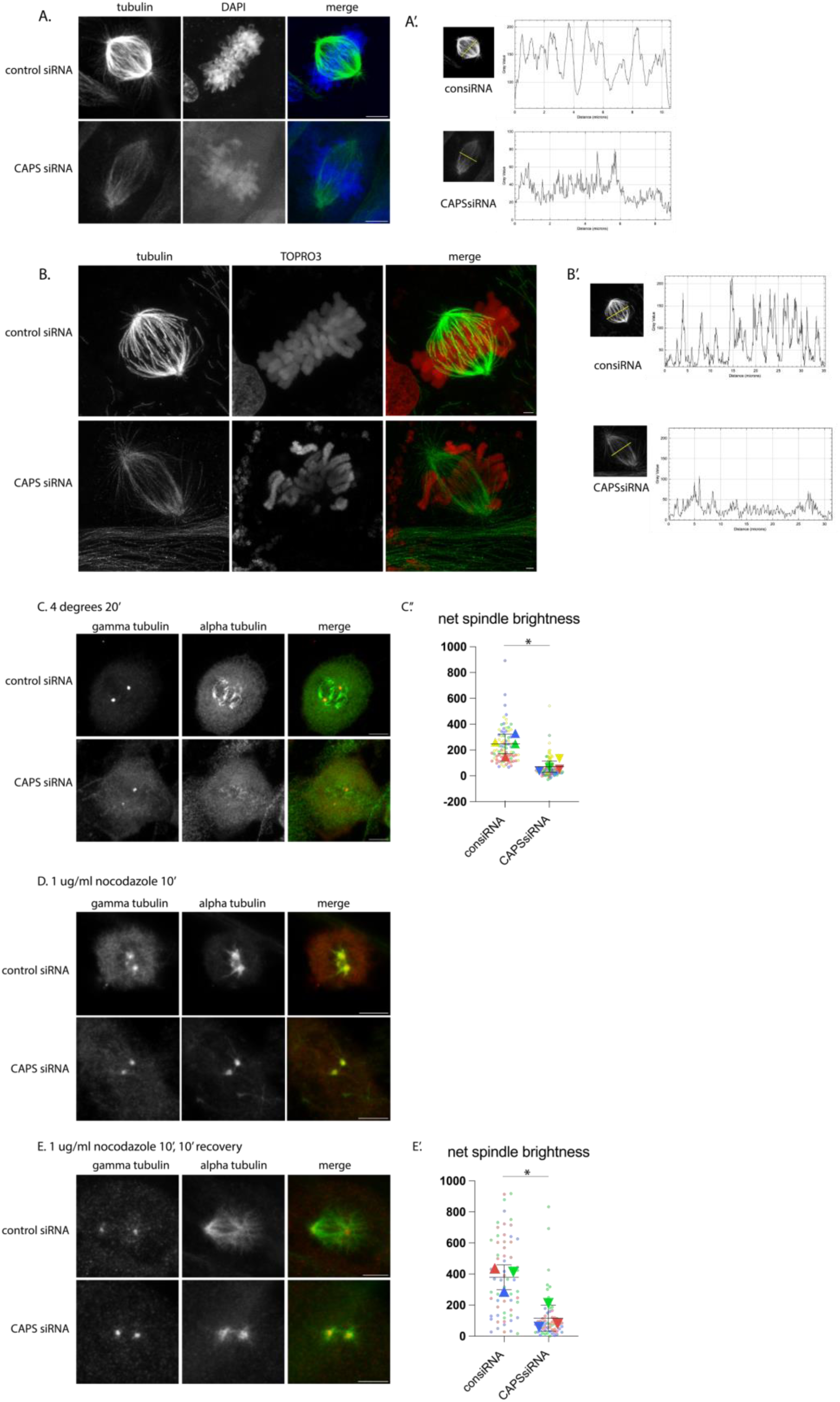
CAPS knockdown reduces the presence of kinetochore fibers. A) Z-stack max projection shows that kinetochore fibers appear reduced in CAPS knockdown cells vs. control knockdown. Densitometry tracing of the spindles (A’) depicts the presence of bundles that correspond to k-fibers; these are reduced in CAPS siRNA-transfected cells. Alpha-tubulin (green), DAPI (blue). B and B’: Expansion microscopy was used to obtain better resolution of k-fibers. Partial z-stack max projection shows that k-fibers are visible in control siRNA but are lacking in CAPS siRNA spindles. Alpha-tubulin (green); Topro3 (red) shows DNA. C) Cold stability assay; tubulin staining after 20” at 4 degrees. “Sum slices” Z-stack projection shows control cells retain some kinetochore fibers, while k-fibers are absent in CAPS siRNA-treated cells. C’) Super-plot shows quantitation of net spindle brightness in (C) (Lord et al., 2020); see Methods; 15-26 cells evaluated/condition; N=4). D) High levels of nocodazole (1ug/ml) completely depolymerized microtubules in both control and CAPS siRNA-treated cells; “sum slices” z-stack projections. E) 10’ after nocodazole washout, spindle microtubules in control siRNA-treated cells reformed more readily than in CAPS siRNA-treated cells; sum slices z-stack projections. E’) Quantitation of net spindle brightness, as in (C’) (20-22 cells evaluated/condition; N=3).

Kinetochore fibers are more resistant to depolymerization than other microtubules (Rieder, 1981), and cold-stable assays have been used to evaluate the presence of k-fibers in cells (Zhou et al., 2019; Warren et al., 2020). We performed two assays to assess microtubule stability in cells treated with either control or CAPS siRNAs. Incubation of cells at 4 degrees for 20 minutes caused some reduction in spindle fibers in cells treated with control siRNAs, but more complete reduction in cells treated with CAPS siRNAs (Fig. 6C). Quantitation of spindle brightness supports this observation: control cells had a net mean spindle brightness of 241 vs. 66 for CAPSsiRNA-treated cells (p<.01). An alternate test of spindle stability is to treat cells with nocodazole, which depolymerizes microtubules. Incubation with 1 ug/ml nocodazole for ten minutes causes depolymerization of the spindle in both control and CAPSsiRNA-treated cells (Fig. 6D), but after a ten-minute washout, spindles of consiRNA-treated cells recovered more rapidly than those of CAPSsiRNA-treated cells (6E), suggesting that control cells contained factors that helped stabilize microtubules. Quantitation of the net spindle brightness showed an average spindle brightness of 385 for consiRNA vs. 121 for CAPSsiRNA-treated cells (p<.02). Together, these tests support the hypothesis that CAPS promotes spindle stability, consistent with a role for CAPS in k-fiber formation.

### Purified CAPS bundles microtubules in vitro

To test whether CAPS was capable of bundling microtubules in vitro, we polymerized purified, fluorescently-tagged tubulin into short microtubules and observed the ability of CAPS or control proteins to cross-link the individual strands into bundles. As shown in Fig. 7A, the presence of CAPS and calcium allowed microtubules to assemble into bundles; this was not observed in the presence of the control protein BSA and calcium. To further assess whether CAPS was responsible for this bundling, we labeled purified CAPS and BSA with a 568nm tag (Fig. 7B). Incubation with 488nm-labeled microtubules showed association of CAPS with bundled microtubules in the presence of calcium (Fig. 7C); interestingly, CAPS568 was only associated with microtubule bundles, not individual microtubule strands, consistent with the interpretation that it may be crosslinking fibers together. No BSA568 was associated with microtubules in the presence of Ca2+, nor was CAPS568 associated with microtubules in the presence of EGTA. (Fig 7D-E).

**Figure 7.**
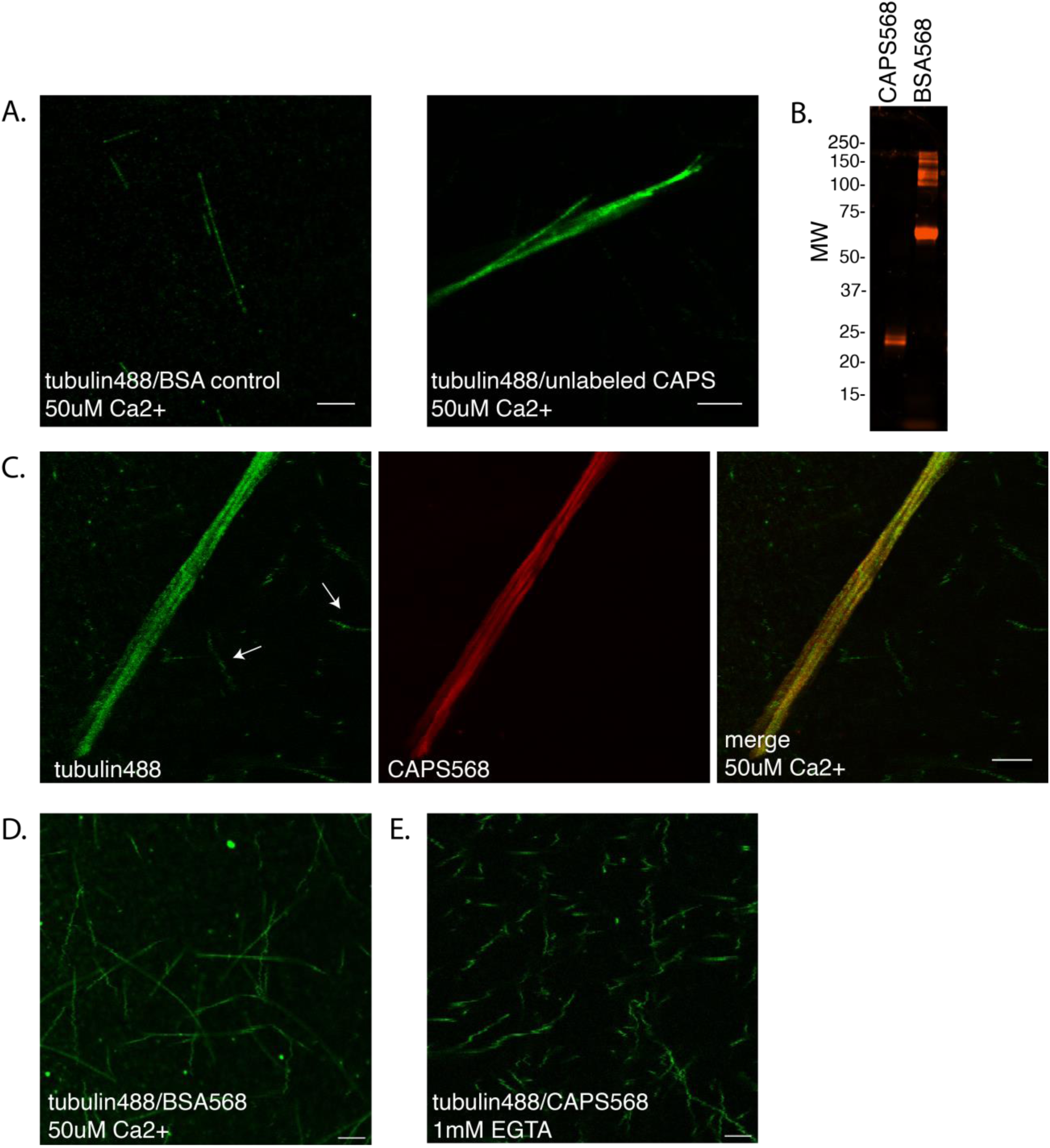
CAPS bundles microtubules in vitro. A) Microtubules were polymerized with 488nm-labeled tubulin and incubated with BSA or CAPS protein in the presence of 50uM Ca2+; bundles only formed in the presence of CAPS. B) SDS-PAGE showing 568nm-tagged CAPS and BSA used in C-E. C-E) CAPS or BSA labeled with a 568-nm fluor was incubated with 488-nm labeled microtubules as in A. C) In the presence of Ca2+, CAPS is associated with bundled microtubules, but not individual microtubules (arrows). D) Labeled BSA did not cause microtubule aggregation, nor did labeled CAPS in the presence of 1mM EGTA (E).

### Overexpression of CAPS does not alter cell proliferation or cell death

Taken together, our results suggest that CAPS is necessary for spindle formation, chromosome congression, and normal progression through mitosis. Given the high level of CAPS expression observed in some human cancers, we also sought to determine if overexpression of CAPS leads to changes in the cell cycle or spindle morphology. N-terminal CAPS-FLAG was transfected into U251 cells and the mitotic index and rate of apoptosis were evaluated. No changes were observed with CAPS overexpression in either rates of mitosis or cell death vs. controls (Fig. 8A, B). To further assess the ability of CAPS to promote proliferation, we also performed an MTT assay of cell growth in low serum media, as was done for HURP overexpression in (Tsou et al., 2003; Prabst et al., 2017). In contrast to HURP, CAPS overexpression did not stimulate cell proliferation more than controls at any serum concentration tested (Fig. 8C). We also analyzed spindle morphology and found no change in spindle length or metaphase plate width, nor in the incidence of uncongressed chromosomes. There was, however, a slight increase in the incidence of multipolar spindles, suggesting that overexpression of CAPS might have some effects on spindle morphology (Supplemental data 3). These results are consistent with the interpretation that CAPS overexpression itself does not alter rates of cell proliferation or cell death. Our data suggest that CAPS is required for spindle integrity and chromosome congression, but that the high level of CAPS observed in many forms of cancer is not a driving force for cell division.

**Figure 8.**
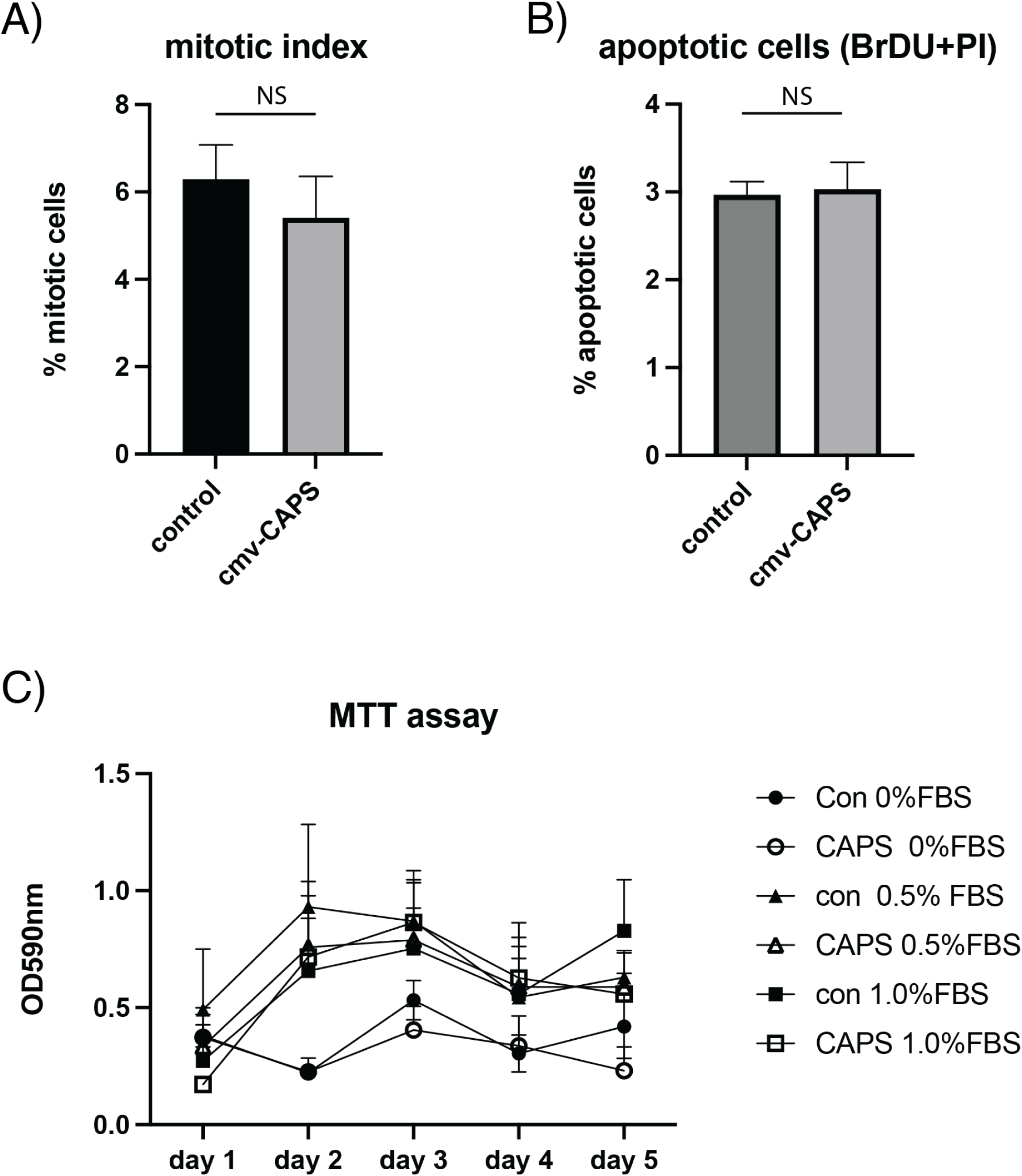
Overexpression of CAPS does not alter cell proliferation or apoptosis. U251MG cells transfected with CAPS-FLAG were evaluated for changes in mitotic index or apoptosis. A) No differences in mitotic index were observed between mock transfected cells and those transfected with CAPS-FLAG B) CAPS-FLAG did not alter the percentage of apoptotic cells as measured by flow cytometry (as in Fig. 5). C) MTT assay shows no significant difference in proliferation between cells transfected with CAPS-FLAG and control (cmv-gfp) at any serum concentration (0%, 0.5%, 1%).

## Discussion

Our results suggest that CAPS is a spindle-associated protein necessary for chromosome congression and progression through mitosis. Several lines of evidence support the hypothesis that CAPS promotes bundling of microtubules to form k-fibers, thus allowing their attachment to chromosomes: CAPS is associated with microtubules of the central spindle, not astral microtubules, nor microtubules in the cytosol. Loss of CAPS appears to reduce k-fiber formation, and knockdown phenotypes are consistent with a lack of attachment between k-fibers and chromosomes. Methods that reduce unbundled microtubules (e.g. cold, nocodazole) appear to affect spindles more in cells lacking CAPS than in cells controls. Finally, purified CAPS appears capable of inducing microtubule bundling in vitro. The dramatic decrease in mitotic index observed with CAPS knockdown could be due to an inability of cells to form a mitotic spindle, though no significant increase in prophase cells was observed. A possible reason for the decrease in mitotic index is that after 48 hours with CAPS knockdown, many cells may have accumulated defects in chromosomal number that might preclude their entry into the cell cycle. The progression of cells through anaphase, even in the presence of multipolar spindles, suggests that genomic integrity would likely be altered after several rounds of cell division.

By what mechanism does CAPS influence microtubule bundling? There are many examples of proteins that dimerize in response to calcium (Donato, 1999; Lee et al., 2017; Ames, 2018), suggesting the presence of calcium could create a CAPS dimer with two tubulin-binding sites. However, experiments by Dong et al., 2008 showed no calcium-induced dimerization of CAPS either by size exclusion chromatography or analytical ultracentrifugation. Alternatively, it is possible that CAPS acts in a complex with other proteins to accomplish microtubule cross-linking. Hepatoma Upregulated Protein (HURP) is a candidate CAPS binding partner that is thought to contribute to k-fiber bundling (Zhang et al., 2018; Huttlin et al, 2021), and we have confirmed coprecipitation of CAPS and HURP in our lab (G.E., *unpublished results*). A model in which CAPS requires a complex of proteins for cross-linking is at odds, however, with our observations of in vitro microtubule bundling in the presence of purified CAPS and tubulin. Since the tubulin used in these experiments is purified from porcine brain, it is possible it may contain some microtubule-associated proteins (MAPS). Alternatively, it is also possible that the high level of purified CAPS used in the experiments was able to overcome low levels of associated proteins.

Bridges of various lengths appear to link the individual microtubules into k-fibers (Booth et al., 2011). Short bridges (14-17nm) include clathrin, TACC3, and ch-TOG, but components of medium (20-27nm) or long (~50nm) bridges have not been identified (Booth et al., 2011; Nixon et al., 2015). It is possible, then that CAPS and possibly associated proteins could be components of the observed medium- or long-bridges. The phenotype observed with CAPS reduction is similar to that seen in other studies that examined the effects of HURP knockdown, e.g. increases in uncongressed chromosomes and multipolar spindles and retention of spindle assembly checkpoint proteins (Wong and Fang, 2006). However, the localization of CAPS on the spindle is not identical to that of HURP: while HURP is concentrated near the kinetochores (Sillje et al., 2006), CAPS is distributed evenly along central spindle microtubules. Further experiments will be necessary to determine the nature of the interactions between CAPS and HURP and whether the two proteins act cooperatively to ensure k-fiber formation and chromosomal attachment.

CAPS has a close homolog in humans called CAPS-like, or CAPSL, that shares 53% sequence identity with CAPS at the amino acid level. The expression level of CAPSL is generally lower than that of CAPS, and its expression pattern is somewhat different (Karlsson et al., 2021; Human Protein Atlas, proteinatlas.org), but it appears to bind to similar proteins, including tubulin and HURP (Huttlin et al., 2021). Thus, it is possible that CAPS and CAPSL perform overlapping functions during spindle formation. Loss of both proteins from cells will need to be tested to determine if this produces a stronger loss of function phenotype.

## Methods

### Antibodies

The following antibodies were used for immunohistochemistry and western blots: mouse anti-tubulin clone B-5-1-2(Millipore Sigma), rabbit anti-CAPS (Abcam EPR15631), mouse anti-HEC1 (Millipore Sigma SAB2702380), mouse anti-FLAG M2 (Sigma-Aldrich), rabbit BubR1 (Proteintech 11504-2AP), rabbit Mad2L1 (AbClone A11469), mouse anti-HistoneH3 (Abcam 14955). Secondary antibodies used for staining were donkey anti-mouse or anti-rabbit Alexa 488 (Invitrogen), donkey anti-rabbit or anti-mouse Cy3 (Jackson Immunoresearch), and donkey anti-rabbit or anti-mouse Alexa 647 (Jackson Immunoresearch).

### Immunohistochemistry

U251-MG or SKBR3 human cells were cultured by standard methods in DME, 10% FBS, Penn/Strep on #1.5 coverslips. Cells were transfected with siRNAs or plasmids using Lipofectamine 3000 according to manufacturer’s instructions. Cells were fixed either with −20 degree methanol for 5 minutes (for anti-tubulin staining), or 4% paraformaldehyde in PBS for 20 minutes (e.g. for gfp or anti-CAPS staining). Cells were incubated with primary antibodies overnight (1:600 anti-tubulin, 1:300 all other antibodies). After staining, cells were mounted in Prolong Diamond with DAPI (Invitrogen) and imaged using a Leica SP8 confocal microscope.

### DNA constructs

The CAPS gene was obtained from DNAsu. CAPS-FLAG was created by Gateway cloning the CAPS gene into pDEST-N-terminal FLAG (Addgene #18700), providing expression of N-terminally FLAG-tagged CAPS control of the cmv-promoter. CAPS-gfp was created by Gateway cloning the CAPS gene into pDEST-CMV-C-EGFP (Addgene #122844), to create an C-terminally-tagged gfp construct under control of the cmv promoter.

### siRNA knockdown and CAPS overexpression

For most experiments, 30pmol/ml CAPS siRNA hs.Ri.CAPS.13.3 (IDT) was transfected into cells using either RNAiMax (Invitrogen) or Lipofectamine 3000 (Invtrogen) with equivalent results. The sequence (GUUUAAUGUAGUUCCCAAAUACATT) corresponds to the 3’UTR of the human CAPS gene; there are no known off-targets. Some experiments were repeated with a second siRNA (hs.Ri.CAPS.13.1; sequence GGAAUAAAAAUAGCAGGUAACAUGT) with similar results. Rescue of the siRNA-induced phenotype was accomplished by co-transfection of CAPS-FLAG along with the siRNA. N-terminal FLAG-tagged CAPS was created by recombining the human CAPS gene (CAPS/pENTR; DNAsu HscD00514950) into pEZFLAG (Addgene#18700; Guo et al., 2008) using Gateway cloning. For CAPS siRNA rescue, 1 ug cmv-CAPS-FLAG was co-transfected with 30pmol hs.Ri.CAPS.13.3 siRNA using Lipofectamine 3000. For all experiments, transfection of 30pmol/ml Mission siRNA Universal Negative Control #1 (Sigma-Aldrich) was used as a control.

### Spin down assay

A CAPS-GST construct was created in pGEX-6-P1 vector using Gateway recombinant cloning; pGEX6P1-DEST-FLAG was a gift from Andrew Jackson & Martin Reijns (Addgene plasmid # 119754; http://n2t.net/addgene:119754; RRID:Addgene_119754).CAPS-GST protein was expressed in BL21Star bacteria (NEB); the protein was isolated by binding to glutathione-Sepharose (Pierce) and eluted from the column by incubation with either reduced glutathione (to retain the GST tag) or with 10 units Prescission Protease (Genscript) to obtain protein without a tag. Eluted proteins were checked by SDS-PAGE to reveal single bands of 48 kd (with tag) or 23 kd (without tag). Microtubules were polymerized in vitro from purified tubulin dimers (Cytoskeleton, Inc,; Zeng et al., 2010). To evaluate whether CAPS coprecipitated with polymerized microtubules, purified CAPS-GST protein was mixed with increasing concentrations of microtubules in Ca2+ Binding-Buffer (80 mM PIPEs, 2 mM MgCl2, 50 uM CaCl2, 1 mg/ml BSA, 10 uM leupeptin (Sigma Aldrich L-2884), 1 uM Calpain Inhibitor Peptide (Sigma-Aldrich C-9181;); BSA was included to prevent non-specific binding, while leupeptin and Calpain Inhibitor Peptide were included to reduce activity of calcium-activated proteases. Samples were spun at 100,000g in a Beckman airfuge for 30 minutes to pellet microtubules. To prevent non-specific binding of CAPS protein to the centrifuge tubes, tubes were pre-incubated with 10 mg/ml BSA. After centrifugation, the supernatant was removed and the pellet was washed 3X with Ca2+-Binding buffer. Proteins in the supernatant and pellet were solubilized in 1XLDS sample buffer (Sigma) and analyzed by SDS-PAGE and western blots with anti-CAPS and anti-alpha-tubulin antibodies.

### Assays for microtubule stability

To identify cold-stable k-fibers, cells grown on coverslips were washed once with 4 degree PBS and placed at 4 degrees for 20 minutes, followed by fixation with −20 degree MeOH for three minutes. To assess stability of microtubules in response to nocodazole treatment, cells were incubated with 1ug/ml nocodazole in regular cell culture media for 10 minutes. The nocodazole was then removed by washing the cells twice with room-temperature PBS, and microtubules were allowed to reform in fresh, 37-degree tissue culture media for 10 minutes before fixation with −20 degree MeOH for three minutes. After staining with anti-alpha tubulin and anti-gamma tubulin antibodies, Z-stacks of cells were created using laser scanning microscopy; identical settings were used for control and CAPS knockdown cells. Spindle brightness was quantified as follows: Z-projections were created using SUM SLICES in Image J, then an ellipse was drawn around the spindle area, using the gamma-tubulin staining as a guide. The mean pixel brightness in the ellipse was quantified in ImageJ using the Measure tool, and the background mean pixel brightness was measured in a small ellipse to the right of the spindle. The net brightness was calculated as the spindle brightness minus background. Three independent experiments were quantified; results are presented as super-plots showing individual values as well as mean +/− standard deviation.

### In vitro bundling assay

50ug (1nM) Tubulin dimers were mixed with 10ug (0.2nM) FITC-labeled alpha tubulin (5:1) in 20ul of 6% glycerol, 80mM PIPES, 2 mMMgCl2 and incubated at 37 degrees 20 minutes to form ~5 um long FITC-labeled microtubules (Zhu et al., 2018). MTs were diluted into 200ul either Ca2+- or EGTA-containing buffers containing taxol (80mM PIPES, 2mM MgCl2, 20uM taxol, and either 100uM CaCl2 or 1mM EGTA). Purified CAPS protein (without GST-tag; see above) or BSA (as a control) were fluorescently tagged with a 568nm label using Mix-N-Stain CF 568 kit (Millipore Sigma). Unincorporated label was removed by a size filtration spin column and incorporation of the tag was checked by SDS-PAGE. Microtubules (approximately 20uM tubulin) were incubated with 4.5uM labeled or unlabeled CAPS protein or BSA for 20 minutes and visualized as in (Zhu et al., 2018) using a Leica SP8 confocal microscope. <u>Apoptotic assays:</u> Apoptotic cells were identified by labeling 3’ ends of DNA using a BrdU apoptosis kit (Fisher Scientific cat. 88-6671-88) and anti-BRDU antibodies, followed by flow cytometry on a BD Accuri C6 plus flow cytometer. Alternatively, cells were fixed and stained with antibodies against cleaved Caspase-3 (Cell Signaling, D175, clone 5A1E) and propidium iodide and evaluated using flow cytometry.

### MTT assay

The MTT assay for cell proliferation was performed using MTT Cell Growth Assay Kit (Sigma Aldrich CT02). HEK293 cells transfected with either cmv-gfp (control; Addgene 11153) or cmv-CAPS-FLAG were plated in a 96-well plate, with 10^4^ cells/well. Cells were incubated (in quadruplicate) with Dulbeccos Modified Eagles media supplemented with 0% fetal bovine serum (FBS), 0.5% FBS, or 1.0% FBS. For 5 days, at each 24 hr timepoint, cells were incubated with 100 ul MTT reagent for 3 hrs, then lysed with 150ul DMSO. After 15 minutes, absorbance at 590nm was determined in a plate reader (Biotek).

### Contributions

B.S. performed phenotypic characterization and flow cytometry experiments and edited the manuscript, Q.N. created CAPS constructs and did initial immuneprecipitations to show tubulin binding, S.S. performed staining experiments, some phenotypic characterization, and wrote the manuscript, G.E. cloned the CAPS-gfp construct, showed localization of CAPS-gfp, and edited manuscript.

## Supplemental Figure legends

**Supplemental Figure S1:**
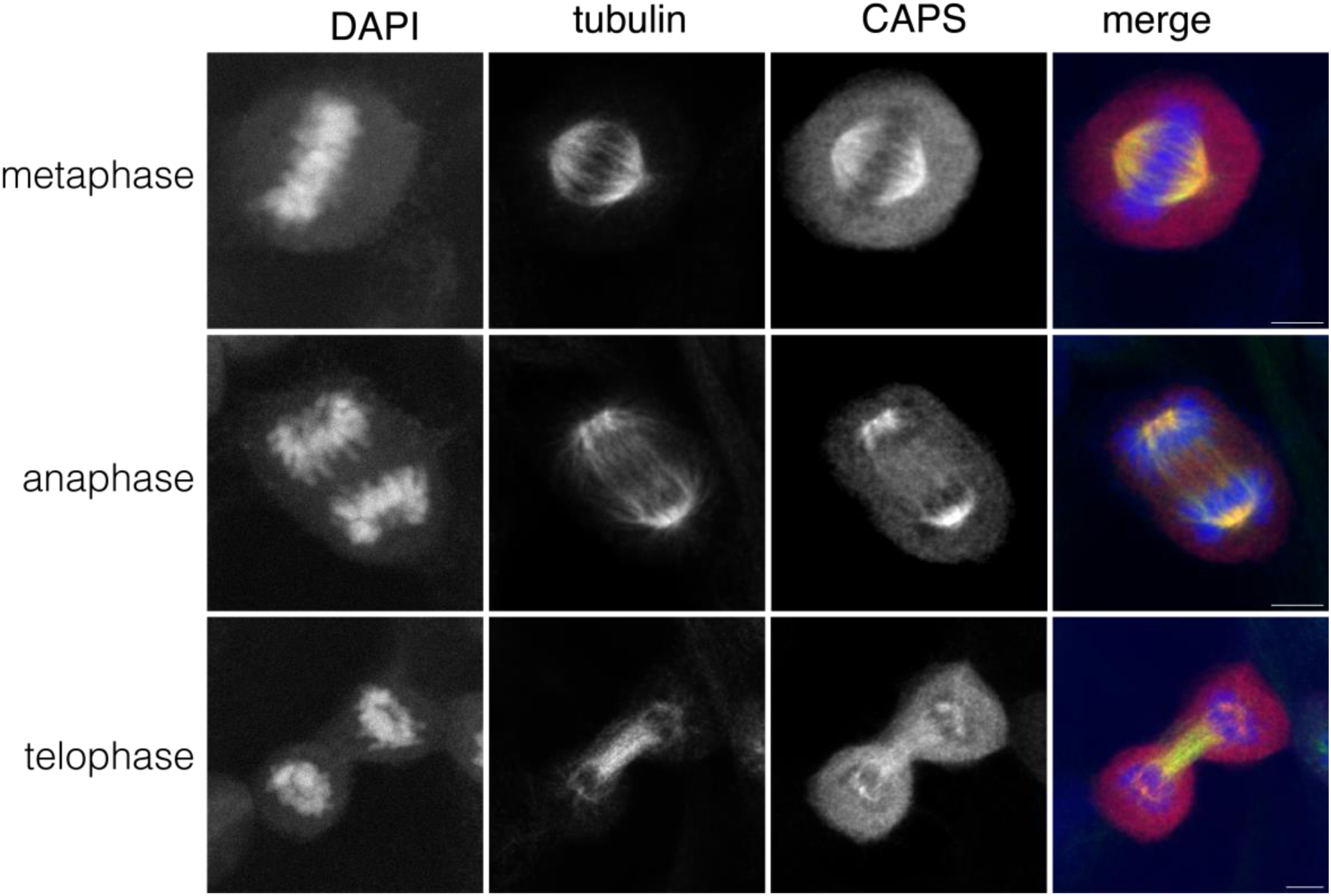
U251MG cells were stained with antibodies against CAPS (red) and alpha-tubulin (green), along with DAPI (blue). Staining shows localization of CAPS on the mitotic spindle, similar to SK-BR3 cells shown in figure 2.

**Supplemental movies S2a and S2b:**

U251MG cells transfected with CAPS siRNA show frequent uncongressed chromosomes and multipolar spindles. Nevertheless, cells often proceed through anaphase, giving rise to three or more daughter cells.

Supplemental 2: (see attached video files)

**Supplemental Figure S3:**
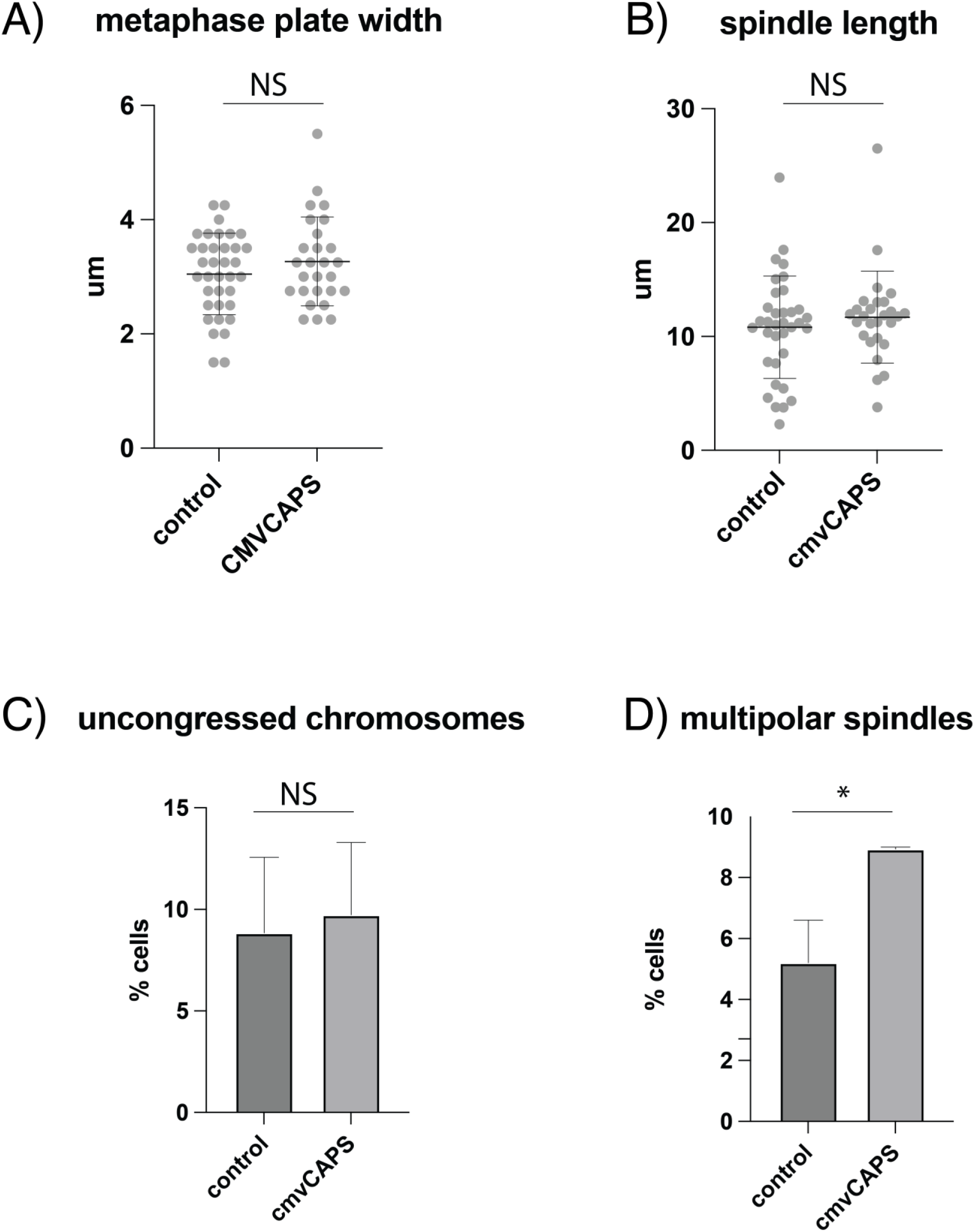
Overexpression of CAPS-FLAG in U251MG cells has little effect on spindle formation. A) metaphase plate width, B) spindle length, and C) the percentage of cells with uncongressed chromosomes are similar between controls and cells overexpressing CAPS-FLAG. D) Overexpression of CAPS-FLAG does, however, produce a slight increase in the percentage of cells with multipolar spindles (p=.01).

## References

Ames, J.B. 2018. Dimerization of Neuronal Calcium Sensor Proteins. Front Mol Neurosci. 11:397.

Andrews, C., Y. Xu, M. Kirberger, and J.J. Yang. 2020. Structural Aspects and Prediction of Calmodulin-Binding Proteins. International journal of molecular sciences. 22.

Booth, D.G., F.E. Hood, I.A. Prior, and S.J. Royle. 2011. A TACC3/ch-TOG/clathrin complex stabilises kinetochore fibres by inter-microtubule bridging. EMBO J. 30:906–919.

Brouhard, G.J., and L.M. Rice. 2018. Microtubule dynamics: an interplay of biochemistry and mechanics. Nat Rev Mol Cell Biol. 19:451–463.

de Bont, J.M., M.L. den Boer, J.M. Kros, M.M. Passier, R.E. Reddingius, P.A. Smitt, T.M. Luider, and R. Pieters. 2007. Identification of novel biomarkers in pediatric primitive neuroectodermal tumors and ependymomas by proteome-wide analysis. J Neuropathol Exp Neurol. 66:505–516.

Donato, R. 1999. Functional roles of S100 proteins, calcium-binding proteins of the EF-hand type. Biochim Biophys Acta. 1450:191–231.

Dong, H., X. Li, Z. Lou, X. Xu, D. Su, X. Zhou, W. Zhou, M. Bartlam, and Z. Rao. 2008. Crystal-structure and biochemical characterization of recombinant human calcyphosine delineates a novel EF-hand-containing protein family. J Mol Biol. 383:455–464.

El Housni, H., R. Lecocq, and D. Christophe. 1997. Production of dog calcyphosine in bacteria and lack of phosphorylation by the catalytic subunit of protein kinase A in vitro. Mol Cell Endocrinol. 135:93–97.

Erent, M., S. Pagakis, J.P. Browne, and P. Bayley. 1999. Association of calmodulin with cytoskeletal structures at different stages of HeLa cell division, visualized by a calmodulin-EGFP fusion protein. Mol Cell Biol Res Commun. 1:209–215.

Furuta, A., M. Tanaka, W. Omata, M. Nagasawa, I. Kojima, and H. Shibata. 2009. Microtubule disruption with BAPTA and dimethyl BAPTA by a calcium chelation-independent mechanism in 3T3-L1 adipocytes. Endocr J. 56:235–243.

Guo, F., M.Y. Chiang, Y. Wang, and Y.Z. Zhang. 2008. An in vitro recombination method to convert restriction- and ligation-independent expression vectors. Biotechnol J. 3:370–377.

Helassa, N., C. Nugues, D. Rajamanoharan, R.D. Burgoyne, and L.P. Haynes. 2019. A centrosome-localized calcium signal is essential for mammalian cell mitosis. FASEB J. 33:14602–14610.

Huttlin, E.L., R.J. Bruckner, J. Navarrete-Perea, J.R. Cannon, K. Baltier, F. Gebreab, M.P. Gygi, A. Thornock, G. Zarraga, S. Tam, J. Szpyt, B.M. Gassaway, A. Panov, H. Parzen, S. Fu, A. Golbazi, E. Maenpaa, K. Stricker, S. Guha Thakurta, T. Zhang, R. Rad, J. Pan, D.P. Nusinow, J.A. Paulo, D.K. Schweppe, L.P. Vaites, J.W. Harper, and S.P. Gygi. 2021. Dual proteome-scale networks reveal cell-specific remodeling of the human interactome. Cell. 184:3022–3040 e3028.

Izant, J.G. 1983. The role of calcium ions during mitosis. Calcium participates in the anaphase trigger. Chromosoma. 88:1–10.

Kajtez, J., A. Solomatina, M. Novak, B. Polak, K. Vukusic, J. Rudiger, G. Cojoc, A. Milas, I. Sumanovac Sestak, P. Risteski, F. Tavano, A.H. Klemm, E. Roscioli, J. Welburn, D. Cimini, M. Gluncic, N. Pavin, and I.M. Tolic. 2016. Overlap microtubules link sister k-fibres and balance the forces on bi-oriented kinetochores. Nat Commun. 7:10298.

Karlsson, M., C. Zhang, L. Mear, W. Zhong, A. Digre, B. Katona, E. Sjostedt, L. Butler, J. Odeberg, P. Dusart, F. Edfors, P. Oksvold, K. von Feilitzen, M. Zwahlen, M. Arif, O. Altay, X. Li, M. Ozcan, A. Mardinoglu, L. Fagerberg, J. Mulder, Y. Luo, F. Ponten, M. Uhlen, and C. Lindskog. 2021. A single-cell type transcriptomics map of human tissues. Science advances. 7.

Kiewisz, R., G. Fabig, W. Conway, D. Baum, D. Needleman, and T. Muller-Reichert. 2022. Three-dimensional structure of kinetochore-fibers in human mitotic spindles. eLife. 11.

Kletter, T., S. Reusch, T. Cavazza, N. Dempewolf, C. Tischer, and S. Reber. 2022. Volumetric morphometry reveals spindle width as the best predictor of mammalian spindle scaling. J Cell Biol. 221.

Koffa, M.D., C.M. Casanova, R. Santarella, T. Kocher, M. Wilm, and I.W. Mattaj. 2006. HURP is part of a Ran-dependent complex involved in spindle formation. Curr Biol. 16:743–754.

Lawrence, E.J., S. Chatterjee, and M. Zanic. 2023. More is different: Reconstituting complexity in microtubule regulation. J Biol Chem. 299:105398.

Lecocq, R., F. Lamy, and J.E. Dumont. 1979. Pattern of protein phosphorylation in intact stimulated cells: thyrotropin and dog thyroid. Eur J Biochem. 102:147–152.

Lecocq, R., F. Lamy, and J.E. Dumont. 1990. Use of two-dimensional gel electrophoresis and autoradiography as a tool in cell biology: the example of the thyroid and the liver. Electrophoresis. 11:200–212.

Lee, J.J., S.Y. Yang, J. Park, J.E. Ferrell, Jr., D.H. Shin, and K.J. Lee. 2017. Calcium Ion Induced Structural Changes Promote Dimerization of Secretagogin, Which Is Required for Its Insulin Secretory Function. Scientific reports. 7:6976.

Li, F., D. Zhu, Y. Yang, K. Wu, and S. Zhao. 2017. Overexpression of calcyphosine is associated with poor prognosis in esophageal squamous cell carcinoma. Oncology letters. 14:6231–6237.

Li, G., and J.K. Moore. 2020. Microtubule dynamics at low temperature: evidence that tubulin recycling limits assembly. Mol Biol Cell. 31:1154–1166.

Li, Z., W. Min, C. Huang, S. Bai, M. Tang, and X. Zhao. 2010. Proteomics-based approach identified differentially expressed proteins with potential roles in endometrial carcinoma. Int J Gynecol Cancer. 20:9–15.

Lord, S.J., K.B. Velle, R.D. Mullins, and L.K. Fritz-Laylin. 2020. SuperPlots: Communicating reproducibility and variability in cell biology. J Cell Biol. 219.

Maiato, H., and E. Logarinho. 2014. Mitotic spindle multipolarity without centrosome amplification. Nat Cell Biol. 16:386–394.

Matkovic, J., S. Ghosh, M. Cosic, S. Eibes, M. Barisic, N. Pavin, and I.M. Tolic. 2022. Kinetochore- and chromosome-driven transition of microtubules into bundles promotes spindle assembly. Nat Commun. 13:7307.

McEwen, B.F., A.B. Heagle, G.O. Cassels, K.F. Buttle, and C.L. Rieder. 1997. Kinetochore fiber maturation in PtK1 cells and its implications for the mechanisms of chromosome congression and anaphase onset. J Cell Biol. 137:1567–1580.

Moisoi, N., M. Erent, S. Whyte, S. Martin, and P.M. Bayley. 2002. Calmodulin-containing substructures of the centrosomal matrix released by microtubule perturbation. J Cell Sci. 115:2367–2379.

Nixon, F.M., C. Gutierrez-Caballero, F.E. Hood, D.G. Booth, I.A. Prior, and S.J. Royle. 2015. The mesh is a network of microtubule connectors that stabilizes individual kinetochore fibers of the mitotic spindle. eLife. 4.

O’Toole, E., M. Morphew, and J.R. McIntosh. 2020. Electron tomography reveals aspects of spindle structure important for mechanical stability at metaphase. Mol Biol Cell. 31:184–195.

Oyama, Y., T. Shishibori, K. Yamashita, T. Naya, S. Nakagiri, H. Maeta, and R. Kobayashi. 1997. Two distinct anti-allergic drugs, amlexanox and cromolyn, bind to the same kinds of calcium binding proteins, except calmodulin, in bovine lung extract. Biochem Biophys Res Commun. 240:341–347.

Phengchat, R., H. Takata, S. Uchiyama, and K. Fukui. 2017. Calcium depletion destabilises kinetochore fibres by the removal of CENP-F from the kinetochore. Scientific reports. 7:7335.

Polak, B., P. Risteski, S. Lesjak, and I.M. Tolic. 2017. PRC1-labeled microtubule bundles and kinetochore pairs show one-to-one association in metaphase. EMBO Rep. 18:217–230.

Prabst, K., H. Engelhardt, S. Ringgeler, and H. Hubner. 2017. Basic Colorimetric Proliferation Assays: MTT, WST, and Resazurin. Methods Mol Biol. 1601:1–17.

Rieder, C.L. 1981. The structure of the cold-stable kinetochore fiber in metaphase PtK1 cells. Chromosoma. 84:145–158.

Rodrigues, M.A., D.A. Gomes, M.F. Leite, W. Grant, L. Zhang, W. Lam, Y.C. Cheng, A.M. Bennett, and M.H. Nathanson. 2007. Nucleoplasmic calcium is required for cell proliferation. J Biol Chem. 282:17061–17068.

Saoudi, Y., B. Rousseau, J. Doussiere, S. Charrasse, C. Gauthier-Rouviere, N. Morin, C. Sautet-Laugier, E. Denarier, R. Scaife, C. Mioskowski, and D. Job. 2004. Calcium-independent cytoskeleton disassembly induced by BAPTA. Eur J Biochem. 271:3255–3264.

Shao, W., Q. Wang, F. Wang, Y. Jiang, M. Xu, and J. Xu. 2016. Abnormal expression of calcyphosine is associated with poor prognosis and cell biology function in colorectal cancer. OncoTargets and therapy. 9:477–487.

Sillje, H.H., S. Nagel, R. Korner, and E.A. Nigg. 2006. HURP is a Ran-importin beta-regulated protein that stabilizes kinetochore microtubules in the vicinity of chromosomes. Curr Biol. 16:731–742.

Simunic, J., and I.M. Tolic. 2016. Mitotic Spindle Assembly: Building the Bridge between Sister K-Fibers. Trends Biochem Sci. 41:824–833.

Subramanian, R., E.M. Wilson-Kubalek, C.P. Arthur, M.J. Bick, E.A. Campbell, S.A. Darst, R.A. Milligan, and T.M. Kapoor. 2010. Insights into antiparallel microtubule crosslinking by PRC1, a conserved nonmotor microtubule binding protein. Cell. 142:433–443.

Tanaka, T.U., and A. Desai. 2008. Kinetochore-microtubule interactions: the means to the end. Curr Opin Cell Biol. 20:53–63.

Tombes, R.M., and G.G. Borisy. 1989. Intracellular free calcium and mitosis in mammalian cells: anaphase onset is calcium modulated, but is not triggered by a brief transient. J Cell Biol. 109:627–636.

Tsou, A.P., C.W. Yang, C.Y. Huang, R.C. Yu, Y.C. Lee, C.W. Chang, B.R. Chen, Y.F. Chung, M.J. Fann, C.W. Chi, J.H. Chiu, and C.K. Chou. 2003. Identification of a novel cell cycle regulated gene, HURP, overexpressed in human hepatocellular carcinoma. Oncogene. 22:298–307.

Valdez, V.A., L. Neahring, S. Petry, and S. Dumont. 2023. Mechanisms underlying spindle assembly and robustness. Nat Rev Mol Cell Biol. 24:523–542.

Warren, J.D., B. Orr, and D.A. Compton. 2020. A comparative analysis of methods to measure kinetochore-microtubule attachment stability. Methods Cell Biol. 158:91–116.

Wong, J., and G. Fang. 2006. HURP controls spindle dynamics to promote proper interkinetochore tension and efficient kinetochore capture. J Cell Biol. 173:879–891.

Xu, N., K.Q. Luo, and D.C. Chang. 2003. Ca2+ signal blockers can inhibit M/A transition in mammalian cells by interfering with the spindle checkpoint. Biochem Biophys Res Commun. 306:737–745.

Yuasa, H.J., A. Nakatomi, T. Suzuki, and M. Yazawa. 2002. Genomic structure of the sponge, Halichondria okadai calcyphosine gene. Gene. 298:21–27.

Zeng, S.X., Y. Li, Y. Jin, Q. Zhang, D.M. Keller, C.M. McQuaw, E. Barklis, S. Stone, M. Hoatlin, Y. Zhao, and H. Lu. 2010. Structure-specific recognition protein 1 facilitates microtubule growth and bundling required for mitosis. Mol Cell Biol. 30:935–947.

Zhang, Y., L. Tan, Q. Yang, C. Li, and Y.C. Liou. 2018. The microtubule-associated protein HURP recruits the centrosomal protein TACC3 to regulate K-fiber formation and support chromosome congression. J Biol Chem. 293:15733–15747.

Zhou, H., T. Zheng, T. Wang, Q. Li, F. Wang, X. Liang, J. Chen, and J. Teng. 2019. CCDC74A/B are K-fiber crosslinkers required for chromosomal alignment. BMC Biol. 17:73.

Zhu, Y., W. Tan, and W.L. Lee. 2018. An in vitro Microscopy-based Assay for Microtubule-binding and Microtubule-crosslinking by Budding Yeast Microtubule-associated Protein. Bio Protoc. 8.

